# Mode selection mechanism in traveling and standing waves revealed by Min wave reconstituted in artificial cells

**DOI:** 10.1101/2022.01.10.475761

**Authors:** Sakura Takada, Natsuhiko Yoshinaga, Nobuhide Doi, Kei Fujiwara

## Abstract

Reaction-diffusion coupling (RDc) generates spatiotemporal patterns, including two dynamic wave modes: traveling and standing waves. Although mode selection plays a significant role in the spatiotemporal organization of living cell molecules, the mechanism for selecting each wave mode remains elusive. Here, we investigated a wave mode selection mechanism using Min waves reconstituted in artificial cells, emerged by the RDc of MinD and MinE. Our experiments and theoretical analysis revealed that the balance of membrane binding and dissociation from the membrane of MinD determines the mode selection of the Min wave. We successfully demonstrated that the transition of the wave modes can be regulated by controlling this balance and found hysteresis characteristics in the wave mode transition. These findings highlight a novel role of the balance between activators and inhibitors as a determinant of the mode selection of waves by RDc and depict a novel mechanism in intracellular spatiotemporal pattern formations.

**Teaser:** Activator-inhibitor balance determines whether a nonlinear wave in live cells becomes a traveling wave or standing wave.

## Introduction

Spatiotemporal patterns of molecules in living cells are formed by the self-organization of molecules. A striking mechanism that induces this self-organization is reaction-diffusion coupling (RDc). RDc induces two types of self-assembly: static and dynamic patterns. Examples of RDc static patterns in living cells are PAR protein patterns, which determine the anterior and posterior poles in the eukaryotic embryo(*1, 2*), and Cdc42 polarization, which contributes to the determination of the budding position of budding yeast(*3, 4*). Dynamic patterns due to RDc are observed as waves of molecular transport. For example, Actin/PIP3/PTEN waves are related to the motility of amoeba cells(*5-8*), the Rho/F-actin wave is related to the cell division of animal cells(*9, 10*), and the Min wave is related to the determination of the cell division plane of bacteria(*11, 12*).

Dynamic patterns caused by RDc are divided into two modes of waves: traveling waves that propagate spatially, and standing waves that oscillate at a fixed point. Both wave modes have been observed in living cells. For example, the actin waves described above are traveling waves(*9, 13*), and the Min wave is considered as a standing wave in living cells(*14, 15*). The modes of the RDc wave are related to its intracellular function. Actin waves guide the direction of cell movement by wave propagation on the membrane(*6, 16*), and the Min wave restricts the initiation of cell division at the cell division plane by enriching the cell division inhibitor at the cell poles through oscillation between the cell poles of the inhibitor(*11, 14*). Therefore, elucidating the mechanism of mode selection between standing and traveling waves is essential for understanding spatiotemporal patterning in cells. Physical picture of the mode selection has been studied in specific systems such as fluid convection of binary mixtures (*17, 18*) and CO oxidization on Pt(*19*). However, the underlying mechanism remains elusive.

Because RDc dynamic waves emerge under nonlinear and far from equilibrium conditions, experiments using defined factors are indispensable for elucidating the mechanism of their spatiotemporal pattern formation. In this regard, the Min system, which uses the Min wave, is a promising material for elucidating the mechanism of wave mode selection for the following two reasons: i) Min waves show both wave modes under certain conditions and ii) the Min wave is the only intracellular RDc wave that can be reconstituted using defined factors in both in vitro (*20-25*)and artificial cells(*26-29*). The Min system consists of MinC, MinD, and MinE, and the RDc of MinDE generates Min waves(*11, 20, 30*). The molecular mechanism is well understood and is shown in Fig. 1A. MinD binds to the membrane in an ATP-dependent manner (*31, 32*)and is recruited to the membrane as a positive feedback loop(*33-35*). MinE, which senses the membrane-binding MinD, binds to the membrane and stimulates the ATPase activity of MinD(*36*). Hydrolyzing ATP bound to MinD triggers the dissociation of MinD from the membrane(*32*). MinE are detached from the membrane after lingering on the membrane for a short time(*24, 33*). Although MinC, which inhibits the polymerization of FtsZ that forms a cytokinetic Z-ring as a cell division initiator(*37*), is not involved in the formation of RDc waves, it co-localizes with MinD to prevent cell division at the cell poles(*11, 14*). Previous studies have shown that MinD, MinE, and ATP are sufficient for the reconstitution of traveling waves on 2D phospholipid membranes(*20*).

**Fig. 1.**
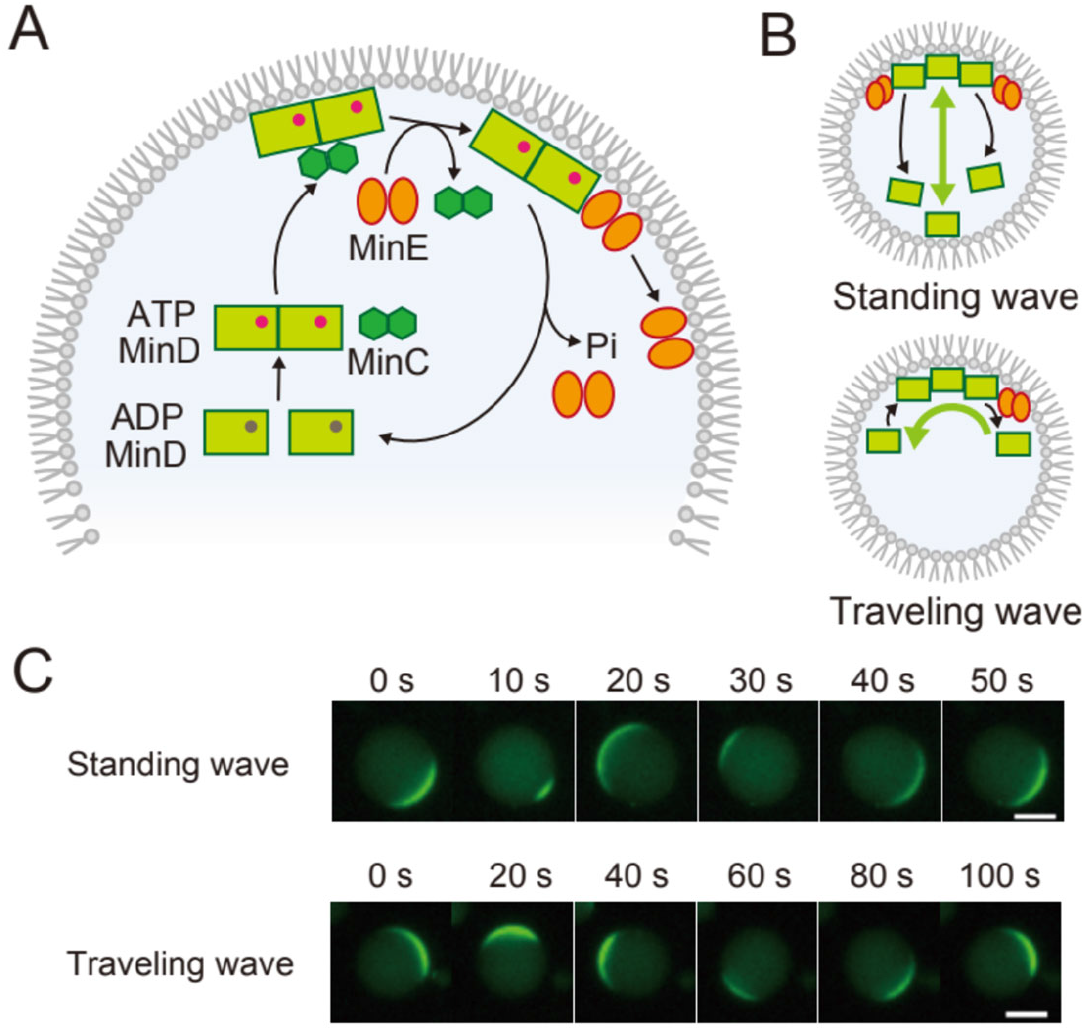
Two dynamic modes of Min waves. (**A**) Molecular mechanism of the Min system. (**B**) The proposed molecular dynamics underlying the traveling and the standing waves. The green arrows indicate the dynamics of Min waves. (**C**) Two modes of Min waves emerged in artificial cells encapsulating 0.1 μM msfGFP-MinC, 1 μM MinD, 1 μM MinE, and 2.5 mM ATP with 100 mg/mL BSA. Scale bars: 10 μm.

The reconstituted system illuminated not only the molecular mechanism, including the effect of the conformational equilibrium of MinE on Min waves(*38, 39*), but also the importance of physicochemical environments, such as salt environments and lipid compositions(*40*). Furthermore, the reconstitution of Min waves in artificial cells revealed that the cell-sized space works as a regulator that determines the emergent conditions and properties of the Min waves(*28*). These cell-sized space effects, derived from closed geometry, the finite-size effect, and the large ratio of the membrane surface to cytosolic bulk volume, indicate that experiments using artificial cells are required to verify the mechanism of the mode selection of dynamic waves.

To date, both standing and traveling waves have been observed in artificial cells (Fig. 1B). For example, a previous study reported that the behaviors of Min proteins spontaneously transit between traveling and standing waves(*27, 28*). As a determinant of wave mode selection, studies using 2D planar membrane chambers that mimic rod-shaped bacteria have proposed that geometry is key to the emergence of standing waves(*22, 23*). However, even in the case of a typical 2D planar membrane, standing waves have also been observed in the case of a MinD mutant(*41*) or MinD depletion(*24*), and both standing waves and traveling waves have been observed in living cells regardless of the geometry(*42, 43*). This suggests that factors other than geometry are also involved in the mode selection of RDc waves.

Here, we investigated the mechanism of mode selection between standing and traveling waves using Min waves reconstituted in artificial cells. We found that the dominant Min wave mode changes depending on the concentration ratio of MinDE and reaction constants, such as the membrane binding affinity of MinE and ATPase activity of MinD. Furthermore, we showed that the transition of the wave mode can be regulated by changing these parameters and successfully recapitulated the results using theoretical analysis. Our results showed that the balance between membrane binding and the dissociation of MinD determines the mode selection of Min waves, implying that the balance between positive and negative feedbacks determines the mode selection of dynamic RDc waves.

## Results

### Dominant Min wave mode changes depending on MinE concentration

Our previous study indicated that the Min wave initially showed a pulsing localization between the membrane and cytosol, transited to the standing wave, and finally settled in the traveling wave(*28*). This Min wave mode transition suggests that the traveling wave is the most stable mode in spherical cells. However, it has been reported that standing waves appear even in spherical cells shaped by the inhibition of the cytoskeleton(*42*). We first assumed that this inconsistency is a consequence of the difference between our reconstitution system and intracellular conditions, such as the concentration ratio of MinD and MinE and/or the fluorescent protein fused to MinDE. MinD and MinE had the same concentration in our reconstituted system(*28*), while the concentration of MinD was 1.4 times that of MinE in living cells(*44*). Fusion of the fluorescent protein to MinDE possibly affects RDc characteristics because of changes in the molecular diffusion rate. In particular, the fusion of fluorescent proteins to the C-terminus of MinE has been suggested to inhibit its function(*45*). Therefore, in this study, we used MinDE without a fluorescent tag, and instead, the Min waves were tracked by a small amount of msfGFP-MinC, which binds to MinD, whose concentration was 1/5^th^ or less of the MinD concentration.

Under the 1 μM MinDE condition, Min waves were found in approximately 90% of the artificial cells, and both standing and traveling waves were observed (Fig. 1C). Because almost all the Min waves observed were traveling waves in our previous study(*28*), the removal of the fluorescent tag was considered as the cause of the appearance of the standing waves. The period of the Min wave was 93 ± 9 s for the traveling wave and 54 ± 8 s for the standing wave, which was approximately half the period of the previous study (# 2 min) (*28*)and closer to the standing wave period in living cells(*14, 46*).

Next, we investigated the effect of the MinDE concentration on the wave modes. MinE concentration was varied in the range of 0.7-2.5 μM while fixing MinD concentration at 1 μM. In this case, MinD localization was tracked using msfGFP-MinC, as described above. Similar to previous studies(*28, 29*), we classified MinC localization into six patterns: cytosol localization (cytosol), homogeneous localization on the membrane (membrane), inhomogeneous patterning on the membrane (inhomogeneous), pulsing between the membrane and the cytosol (pulsing), traveling wave, and standing wave. The ratio of these patterns varied significantly depending on the MinE concentration (Fig. S1). The probability of the appearance of a Min wave decreased as the MinE concentration increased, and MinC was localized in the cytosol for almost all the artificial cells at the MinE concentration of 2.5 μM. However, standing waves became dominant as the concentration of MinE increased for artificial cells with Min waves. A plot of the proportion of standing waves in the artificial cells with Min waves (standing wave/Min wave ratio) showed that the proportion of standing waves was less than 30% at a MinE concentration of 1.5 μM or less, whereas it was more than 60% for MinE concentration over 1.7 μM and almost all Min waves were standing waves at the MinE concentration of 2.2 μM MinE (Fig. 2A, movie S1). These results suggest that the MinE concentration determines the mode selection of the Min wave.

**Fig. 2.**
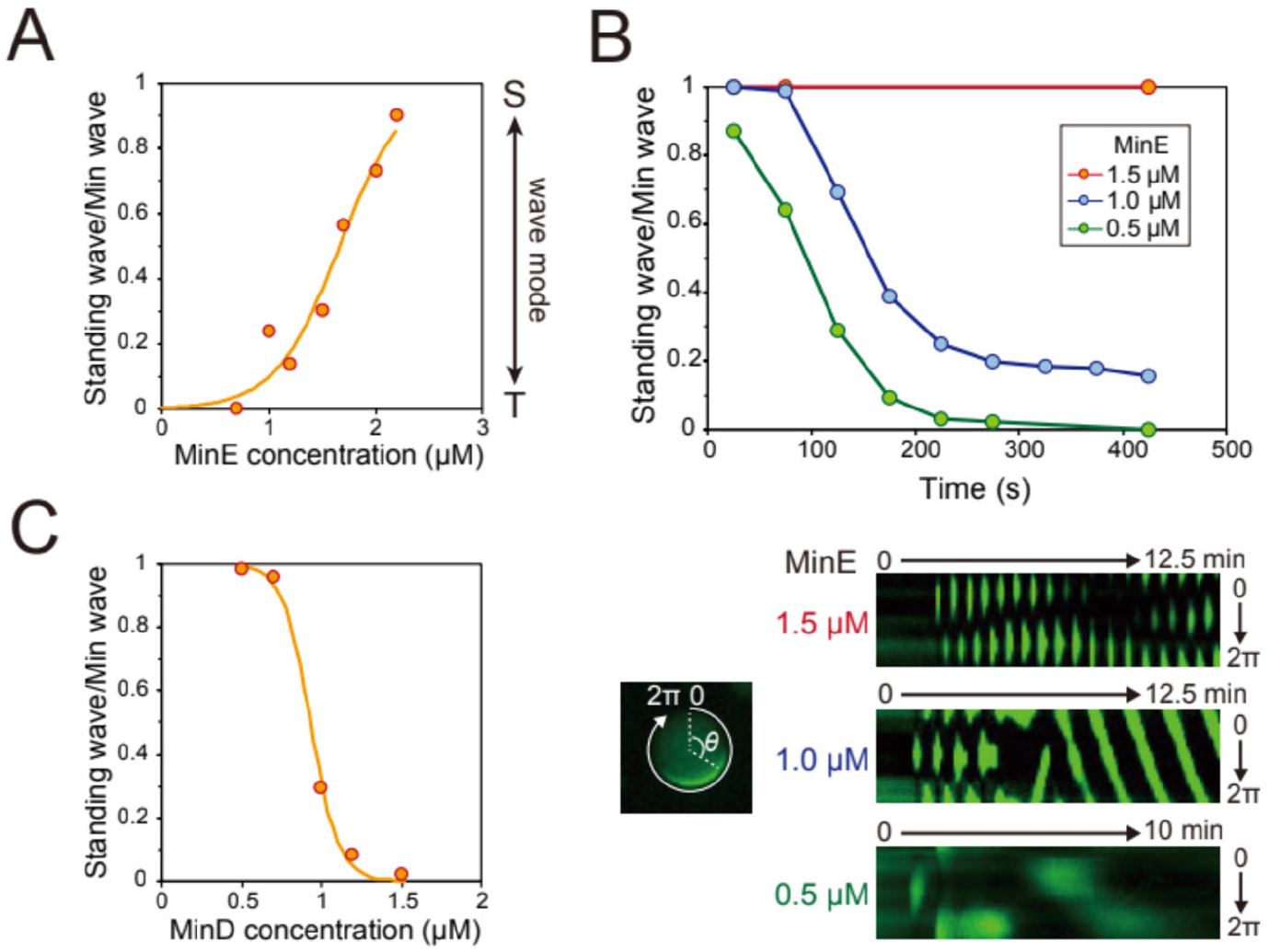
Min protein concentration regulates Min wave modes. (**A-C**) The standing wave/Min wave ratio at various MinE concentrations (**A**), its time development (**B**), and its value at various MinD concentrations (**C**). (**A, C**) The data of the number of artificial cells where Min waves appear are obtained from the observation of more than 150 artificial cells (Figs. S1 and S3). The fitting lines are sigmoidal curves. (**B**) Time development of the standing wave/Min wave ratio at 0.5, 1, and 1.5 μM MinE (top). The data of the artificial cells with Min waves were obtained from the observation of more than 90 artificial cells (Fig. S2). The experiments were performed by encapsulating ADP, creatine kinase, and creatine phosphate instead of ATP. Kymographs of msfGFP-MinC around the membranes of artificial cells at each MinE concentration are shown (bottom).

### Shift in the MinE concentration changes initial transition of Min wave mode after wave emergence

Next, we tested whether the MinE concentration shift affects the transition of the Min wave mode. However, in our first trial, we failed to observe the wave emergence using the procedure from our previous study(*28*). We assumed that this was because the period of the Min wave was shorter than the preparation time of the samples for microscopy. Therefore, we used an ATP regeneration system to generate Min waves. In this experiment, ADP was converted to ATP using a creatine kinase-creatine phosphate system in artificial cells. In the regeneration system, ATP reaches a sufficient concentration for Min wave generation after sample preparation for microscopy. Using this system to observe the initial under the MinE concentrations of 0.5, 1, and 1.5 μM showed that the transition of wave modes was different among the three conditions. Under the MinE concentration of 0.5 μM, the traveling wave appeared just after Min wave emergence in 13% of the artificial cells. In the other artificial cells, the standing wave appeared at the initial stage of the Min wave and quickly transformed into the traveling wave (Fig. 2B and Fig. S2A and Movie S2). Under the MinE concentration of 1 μM, the standing wave appeared at the initial stage of the Min wave in all the artificial cells, and approximately 70 % of the standing waves transformed into traveling waves (Fig. 2B and Fig. S2B, Movie S3). Analysis of kymographs showed that the standing wave transitioned to the traveling wave after approximately four oscillations under the 1 μM MinE condition (Fig. 2B, bottom). The standing wave remained for more than 30 min without transitioning into traveling wave for the MinE concentration of 1.5 μM MinE (Fig. 2B and Fig. S2C and Movie S4). These results showed that the MinE concentration determines the Min wave mode, including the wave mode transition.

### Mode selection of Min wave also depends on MinD concentration

Because the RDc of MinD and MinE causes Min waves, it is plausible that the MinD and MinE concentrations affect the wave mode. Therefore, MinD concentration was varied in the range of 0.5-1.5 μM while MinE concentration was fixed at 1 μM. The dynamics of MinD were tracked using msfGFP-MinC, and Min waves were classified into six patterns as indicated in the MinE experiments. Consequently, the probability of standing waves increased as the MinD concentration decreased in contrast to the MinE case (Fig. 2C and Fig. S3). The proportion of standing waves was 40-60% of the total patterns at low MinD concentrations (0.5, 0.7 μM MinD) and that of the traveling waves was more than 80% at MinD concentrations of 1.2 μM or more (Fig. S3). When the MinD concentration was low, most of the artificial cells without the standing wave showed cytosolic localization (Fig. S3), similar to the condition of high MinE concentration (Fig. S1). Furthermore, the relationship between MinD concentration and the standing wave/Min wave ratio clearly showed that the standing wave was dominant at low MinD concentrations and that the traveling wave was dominant at high MinD concentrations (Fig. 2C and Movie S5). In other words, the mode of the Min wave also depends on the MinD concentration and is not determined by the concentration of either MinD or MinE only but by both concentrations.

### Regulation of the Min wave mode by the activity of Min proteins

Because MinDE concentrations are inversely associated with dominant mode selection, the above results suggest that the concentration ratio of MinDE, rather than their absolute concentration, is important for mode selection. To clarify this point, we focused on the RDc mechanism of the Min system (Fig. 1A). Previous studies have indicated that the membrane binding of MinD(*41*) and MinE(*40*), self-interaction of MinD(*35*), and MinDE dissociation from the membrane via the ATPase activity of the MinDE complex (*47, 48*)are related to concentrations of MinD and MinE. The rates of these reactions were determined by the MinDE concentrations and their reaction constants. Therefore, changes in the reaction constants also act as the concentration shift of MinDE. To verify this hypothesis, we focused on the salt concentration and temperature, which have been shown to affect the membrane binding affinity of MinE and ATPase activity of the MinDE complex, respectively.

First, we investigated the effect of salt concentration on the Min wave mode. The previous study reported that salt concentration changes the membrane binding affinity of MinE by adjusting the electrostatic interaction between the negative charge of the lipids and cationic region of MinE(*40*). Therefore, a high salt concentration was expected to have a similar effect on reducing the MinE concentration (Fig. 3A, left). The concentration of GluK added as salt was varied from 50 to 500 mM, and the wave modes were investigated. Consequently, standing waves became dominant at low salt concentrations, and the percentage of traveling waves increased as the salt concentration increased (Fig. 3A and Fig. S4 and Movie S6). Almost all Min waves were traveling waves at GluK concentrations of 300 mM or more (Fig. 3A). These results clearly indicate that the mode selection of Min waves can be varied by altering the membrane binding affinity of MinE.

**Fig. 3.**
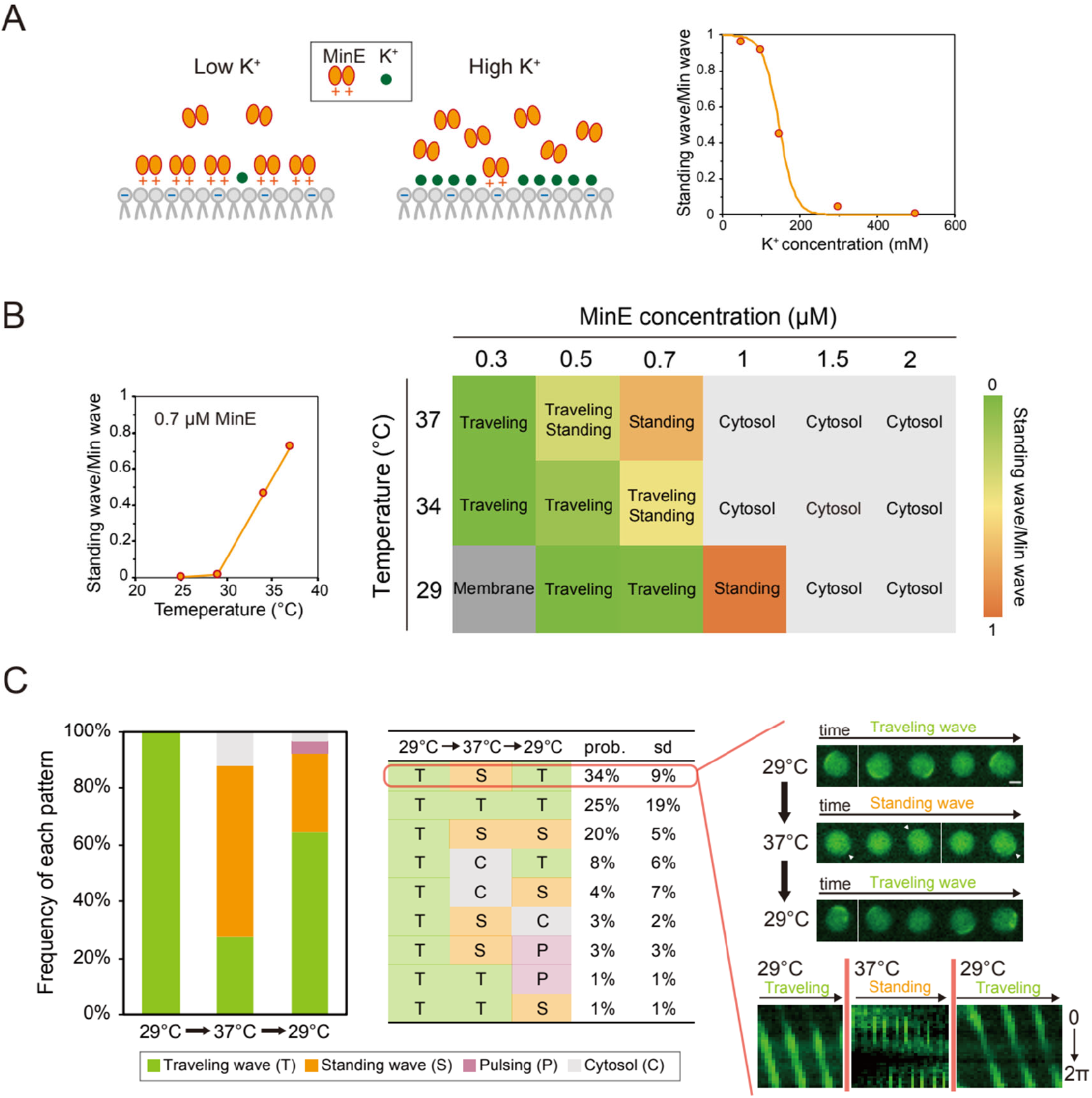
Activities of Min proteins regulate Min wave modes. (**A**) Schematic illustration of the effect of K^+^ concentration on membrane the binding affinity of MinE (left) and changes of the standing wave/Min wave ratio at various K^+^ concentrations (right). The data of the number of artificial cells where Min waves appear were obtained from observing more than 130 artificial cells (fig. S3). The fitting line is a sigmoidal curve. (**B**) Changes in the standing wave/Min wave ratio at various temperatures under a MinE concentration of 0.7 μM (left). Phase diagram of the dynamics of msfGFP-MinC against MinE concentration and the temperature (right). Each square represents the most abundant pattern under each condition. Description of “Traveling/Standing” shows the case of that waves frequently is near a half (0.35-0.65 standing wave/Min wave ratio). The detailed data of all the artificial cells observed are shown in Fig. S5. The color code of the standing wave/Min wave ratio is shown on the right. (**C**) The transition of the frequencies of each pattern induced by temperature shifts. Spatiotemporal patterns of msfGFP-MinC in artificial cells encapsulating 0.2 μM msfGFP-MinC, 1 μM MinD, 0.7 μM MinE, and 2.5 mM ATP with 100 mg/mL BSA are shown on the left(150 artificial cells were counted from three independent experiments.). Only artificial cells where traveling wave emerged at the first temperature (29°C) were counted. The probabilities of each type of spatiotemporal pattern transition in the same artificial cells are shown in the middle (calculated from the same data set as the bar graph. 50 artificial cells, N = 3). Time-lapse images and kymographs of representative pattern transitions are shown on the right. Standing waves are highlighted with white arrows. Time intervals for the time-lapse images are 20 s (traveling waves) and 10 s (standing waves). Scale bar: 20 μm.

Second, we examined whether the ATPase activity of the MinDE complex affects the mode of the Min wave. Although temperature affects various parameters in the RDc of Min waves, previous studies have pointed out that the ATPase activity of the MinDE complex is the most sensitive reaction to temperature(*47, 48*). To verify this, the ATPase activity of the MinDE complex was measured at our typical observation temperature for Min waves (25°C), and at 29 and 37°C. ATPase activity increased as the temperature increased (Fig. S5), and the activation energy was calculated as 7.0 kcal/mol. To verify the effect of increasing the temperature on ATPase activity in the Min wave mode, we varied the Min wave observation temperatures (29, 34, and 37°C). The phase diagram depicted by the dynamics of msfGFP-MinC over the MinE concentration range of 0.3-2 μM showed that an increase in the observation temperature shifts the MinE concentration range that induces the same spatiotemporal pattern (Fig. 3B and Fig. S6). Furthermore, as a result of the calculation of the standing wave/Min wave ratio, the ratio of standing waves increased as the temperature increased at the same MinE concentration (Fig. 3B). In particular, more than 90% of all artificial cells showed standing waves for a MinE concentration of 1 μM MinE at 29°C (Fig. S6). These results demonstrate that the relationship between the MinE concentration and mode of the Min wave shifts depending on the temperature. Even at low MinE concentrations, the behaviors of Min waves under high-temperature conditions were similar to those at high MinE concentrations under low temperature conditions, suggesting that the ATPase activity of the MinDE complex also contributes to the mode selection of the Min wave.

The changes in the dominant mode of the Min wave with temperature indicate that the wave mode can be controlled by shifting the temperature. To verify this point, we observed the effect of the temperature shift on Min wave selection for a MinE concentration of 0.7 μM MinE, where traveling and standing waves are dominant at 29°C and 37°C, respectively (Fig. 3B). As expected, traveling waves were mainly observed at 29°C, and increasing the temperature increased the number of standing waves. We successfully observed the transition by raising the temperature in the same artificial cells (Movie S7). Raising the temperature from 37°C to 29°C transformed 61% of the traveling waves into standing waves (Fig. 3C, left, and Movie S8). It should be noted that this transition is in the opposite direction of the spontaneous transition of the Min wave mode shown in Fig. 2B and observed in our previous study(*28*). Cooling the same artificial cells to 29°C returned the dominant wave mode to the traveling wave mode (Fig. 3C, Movie S8). However, this wave mode return was observed in approximately half of the artificial cells that had standing waves (Fig. 3C, middle), suggesting that the wave mode selection has hysteresis characteristics.

### Hysteresis of Min wave mode analyzed by a protein expression system

Finally, we investigated whether the Min wave mode transition has hysteresis characteristics by MinE concentration shift using a protein expression system. As shown in Fig. 2A, the MinE concentration is a determining factor of the dominant Min wave mode. Therefore, an increase in the MinE concentration in artificial cells was expected to change the dominant wave mode for a Min wave mode transition if the Min waves do not have hysteresis characteristics. To test this hypothesis, MinE was synthesized in artificial cells by co-encapsulating a reconstituted transcription-translation system, the PURE system, and DNA encoding MinE with Min proteins in artificial cells (Fig. 4A).

**Fig. 4.**
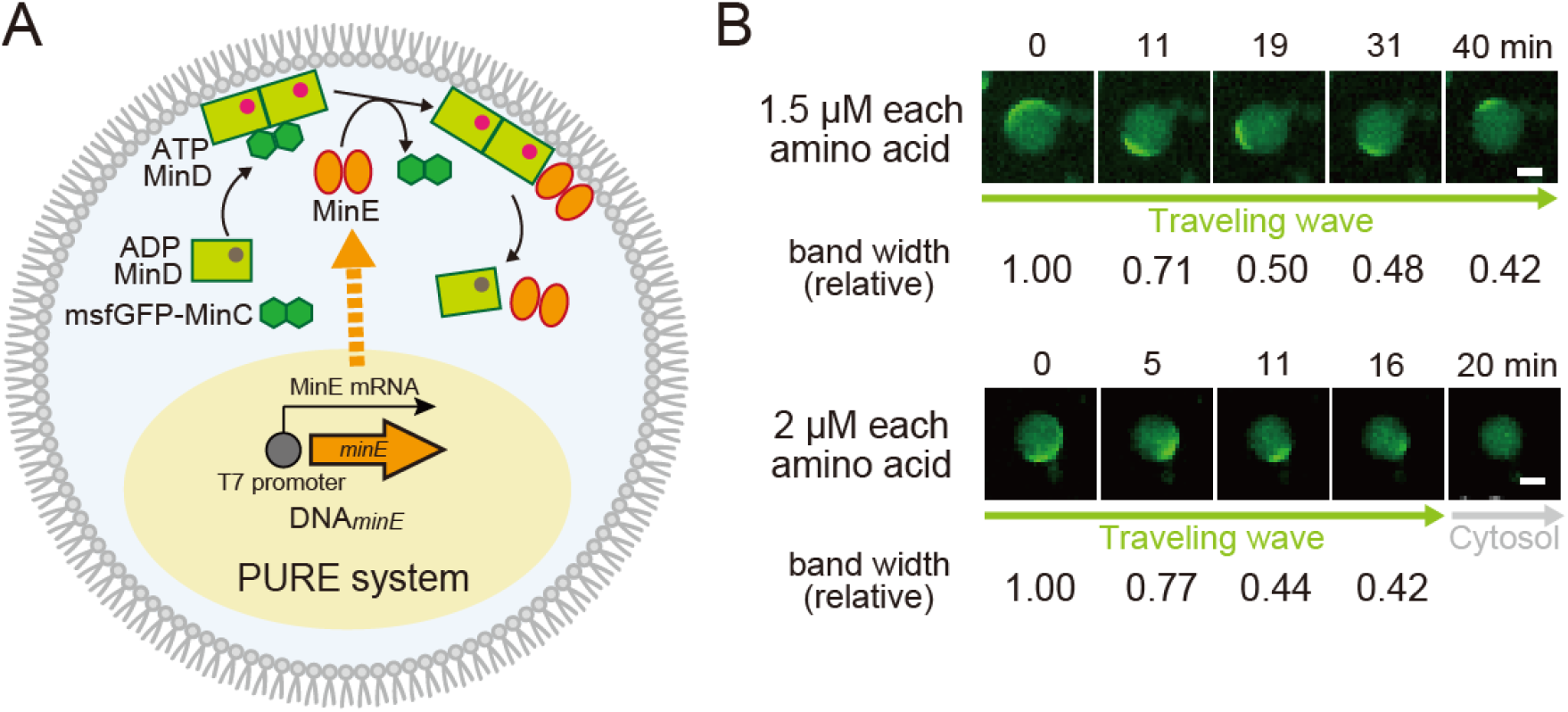
Behaviors of Min waves with MinE synthesis in artificial cells. (**A**) A schematic illustration of the experimental system. In addition to purified msfGFP-MinC, MinD, and MinE, a cell-free transcription and translation system (PURE system) with DNA encoding MinE were encapsulated. (**B**) Time-lapse images of the dynamics of msfGFP-MinC. MinE was synthesized from 1 nM DNA_*minE*_ in artificial cells containing 0.1 μM msfGFP-MinC, 1 μM MinD, 0.7 μM MinE, and amino acids mixtures at 1.5 μM or 2.0 μM each. Relative band widths at each time of the traveling wave normalized by the initial band width are shown. Scale bars: 10 μm.

Because our previous study demonstrated that Min waves reconstituted in artificial cells disappear when an excess amount of MinE is synthesized(*29*), adjusting the MinE synthesis level was necessary. To control protein synthesis levels, we decreased the amount of amino acids required for protein expression. To confirm the reliability of this strategy, NanoLuc, which shows high sensitivity and specificity for quantification, was used as a reporter gene. Consequently, we observed that the concentration of amino acids in the reaction mixture regulated the limited levels of protein synthesis (Fig. S7). Then, we replaced the NanoLuc gene with the MinE gene and analyzed the effect of the increase in the MinE concentration on the Min wave mode. The concentration of the Min proteins contained was set to 1 μM MinD and 0.7 μM MinE, where the traveling wave is dominant. When using amino acids mixtures at 1.25 μM or 1.5 μM each (corresponding to the final MinE concentration synthesized by the protein expression) was supplied, the traveling wave continued and did not transition to a standing wave although the wave width gradually shortened with the synthesis of MinE (Fig. 4B and Movie S9). In contrast, increasing the amino acid concentration to 2 μM each also decreased the wave width in a shorter time, and simultaneously, the traveling wave disappeared after 20 min of the reaction, and the localization of msfGFP-MinC transited to the cytosol (Fig. 4B and Movie S10). The traveling waves did not transition to a standing wave during this process. These results suggest that the mode of the Min wave exhibits hysteresis characteristics.

### Theoretical analysis to capture the relationships between the Min wave mode and parameters

To clarify the mechanism of mode selection between standing and traveling waves, we analyzed the reaction-diffusion equations for the Min system. We used the model that was studied in our previous study (*28*)(see Materials and Methods). The model includes MinD binding on the membrane from the cytosol *ω*_*D*_ and recruitment of MinD *ω*_*dD*_ and MinE *ω*_*E*_ from the cytosol by membrane-bound MinD. In the model, the ADP/ATP exchange *λ* of the MinD in the cytosol after the ATP hydrolysis of MinD (*49*) and the persistent binding of MinE on the membrane (*43*) generate the wave. The latter effect is supplemented by the binding of MinD and MinE *ω*_*ed*_ on the membrane. We also included spontaneous MinE binding on the membrane from the cytosol. This effect is described by a different term *ω*_*eE*_ from the previous study because we are interested in the dependence of wave selection on the concentration of MinE. Spontaneous MinE binding did not change the qualitative features of the phase diagram of the wave states. Without spontaneous MinE binding, the standing waves occur at higher concentrations of MinE, lower concentrations of MinD, and a larger recruitment rate of MinE from the cytosol. Nevertheless, spontaneous MinE binding prevents wave generation at higher concentrations of MinE, resulting in a uniform cytosol state.

Figs. 5A and 5B show the state diagram in the parameter spaces of the MinD and MinE concentrations and the recruitment of MinE(*ω*_*E*_)-MinD(*ω*_*dD*_). When the concentration of MinE was low, MinD accumulated at the membrane and MinE was mainly in the cytosol (Fig. 5C). On the other hand, at higher MinE concentrations, MinD was present in the cytosol. At intermediate concentrations of MinE, both traveling and standing waves appeared depending on the MinDE concentration ratio. When the recruitment of MinE and MinD was varied, the state diagram was qualitatively identical to that of the MinE-MinD concentrations. Standing waves appeared for higher recruitment of MinE and lower recruitment of MinD. In the current model, we added the spontaneous binding of MinE to the membrane. Therefore, waves appear even at *ω*_*E*_ = 0. In both state diagrams, standing waves appeared when MinE frequently attached to the membrane and MinD detached from the membrane, as suggested by the experiments. Conversely, traveling waves dominated when MinD remained attached to the membrane.

**Fig. 5.**
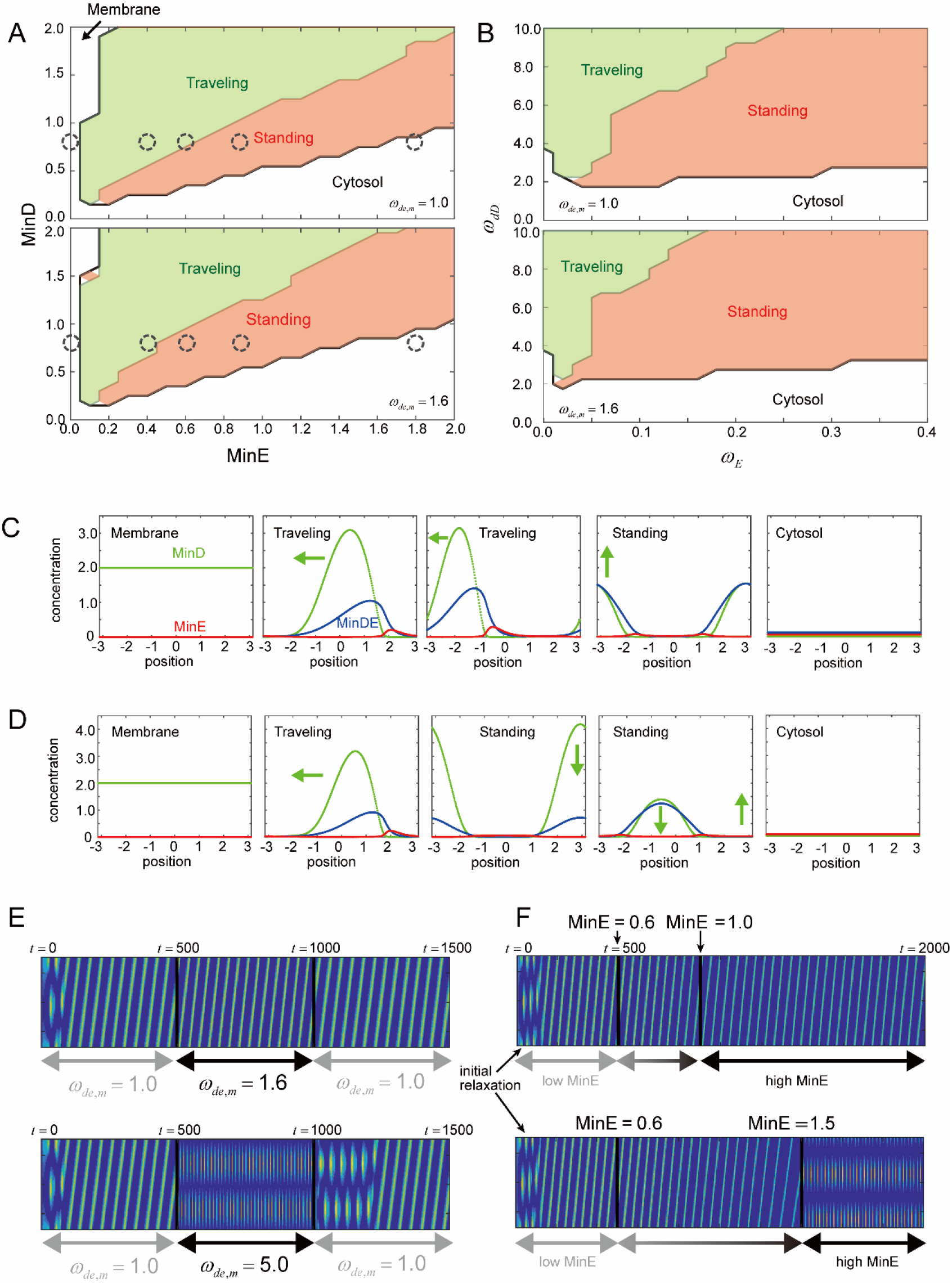
Selection of standing and traveling waves in our theoretical model. (**A, B**) State diagrams of the generated waves under varying MinE-MinD concentrations (**A**), and recruitment of MinE-MinD (**B**). (**C, D**) Spatial distribution of MinD (green), MinE (red), and MinDE (blue) on the membrane at the selected points in the state diagram (**A**) at (**C**) and (**D**). Each distribution from left to right corresponds to the dashed circle in (**A**) from lower to higher concentrations of MinE. (**E, F**) Dynamic change of the waves under varying and the concentration of MinE.

To model the effect of ATPase activity, we used different rates of MinD binding to the membrane *ω*_*de,m*_ (*48*). Following the experimental measurements of ATPase activity at 25 °C and 37 °C (Fig. S5), we set *ω*_*de,m*_ =1 and *ω*_*de,m*_ =1.6, respectively. In both state diagrams of the MinE-MinD concentrations and recruitment, the region where the standing waves occurred broadened with larger ATPase activity. In Fig. 5A, the traveling wave appears at MinD=0.8 and MinE=0.6 when *ω*_*de,m*_ =1 (Fig. 5C). At stronger ATPase activity, *ω*_*de,m*_ =1.6, the standing wave occurred at the same concentrations of MinD and MinE (Fig. 5D). Thus, the model reproduced the same tendency for the selection of the traveling and standing waves.

We also studied the wave transitions under dynamic parameter changes. Following the temperature change and MinE synthesis, we dynamically varied *ω*_*de,m*_ and the concentration of MinE, respectively. We applied a stepwise change in *ω*_*de,m*_ from *t* = 500 to *t* = 1000. After the initial relaxation, the traveling wave switched to the standing wave and returned to the traveling wave at *ω*_*de,m*_ =5.0 (Fig. 5E). Even after *ω*_*de,m*_ was set to the original value, the standing wave remained for some time. This result demonstrates the hysteresis of the transition between the traveling and standing waves. At *ω*_*de,m*_ =1.6, the transition did not occur, although the width of the wave became narrower. Next, we varied the MinE concentration. From *t* = 500, MinE was added to the cytosol at a constant rate (see Materials and Methods). When the final concentration of MinE was 1.0, the system remained in the traveling wave mode (Fig. 5F). On the other hand, when the final concentration was increased to 1.5 (>1.4), the transition to the standing waves occurred (Fig. 5F). Note that in the state diagram in Fig. 5A, a standing wave should appear at MinE > 0.65. Even at a concentration larger than the threshold, the system remained in the traveling wave mode. Together with the experimental result that showed that the traveling wave appears despite a dynamic increase in MinE concentration (Fig. 4B), the results of the theoretical analysis also demonstrate hysteresis of wave mode selection.

## Discussion

Nonlinear waves generated by RDc exhibit two dynamic wave modes: standing and traveling waves. However, the mechanism underlying mode selection has not yet been clarified. Furthermore, the regulation and elucidation mode selection have not been achieved even with Min waves, which show both modes in living cells (*14, 42, 43*)and reconstitution systems(*20, 24, 27, 28*). In our study, using Min waves reconstituted in artificial cells and theoretical analysis, we found that traveling waves were dominant under the conditions of i) a weak inhibitory effect of MinE (shown by MinE (Figs. 2A and 5A)) and salt concentration dependence (Figs. 3A and 5B)) and ii) an increase in the membrane binding rate of MinD (shown by MinD concentration (Figs. 2C and 5A)) (Fig. 6). In contrast, standing waves were dominant under the conditions of i) a strong MinE inhibitory effect (shown by MinE and salt concentration dependence (Figs. 3A and 5B)) and ii) facilitating the membrane dissociation rate of MinD (shown by temperature dependence (Figs. 3B and 5, A and B)) (Fig. 6). Our conclusion is that the balance between the membrane binding and dissociation of MinD, which is determined by the concentration ratio of MinDE and the activities of Min proteins, regulates the Min wave mode (Fig. 6). Our findings imply that the balance between activators and inhibitors is a factor that determines the RDc dynamic wave.

**Fig. 6.**
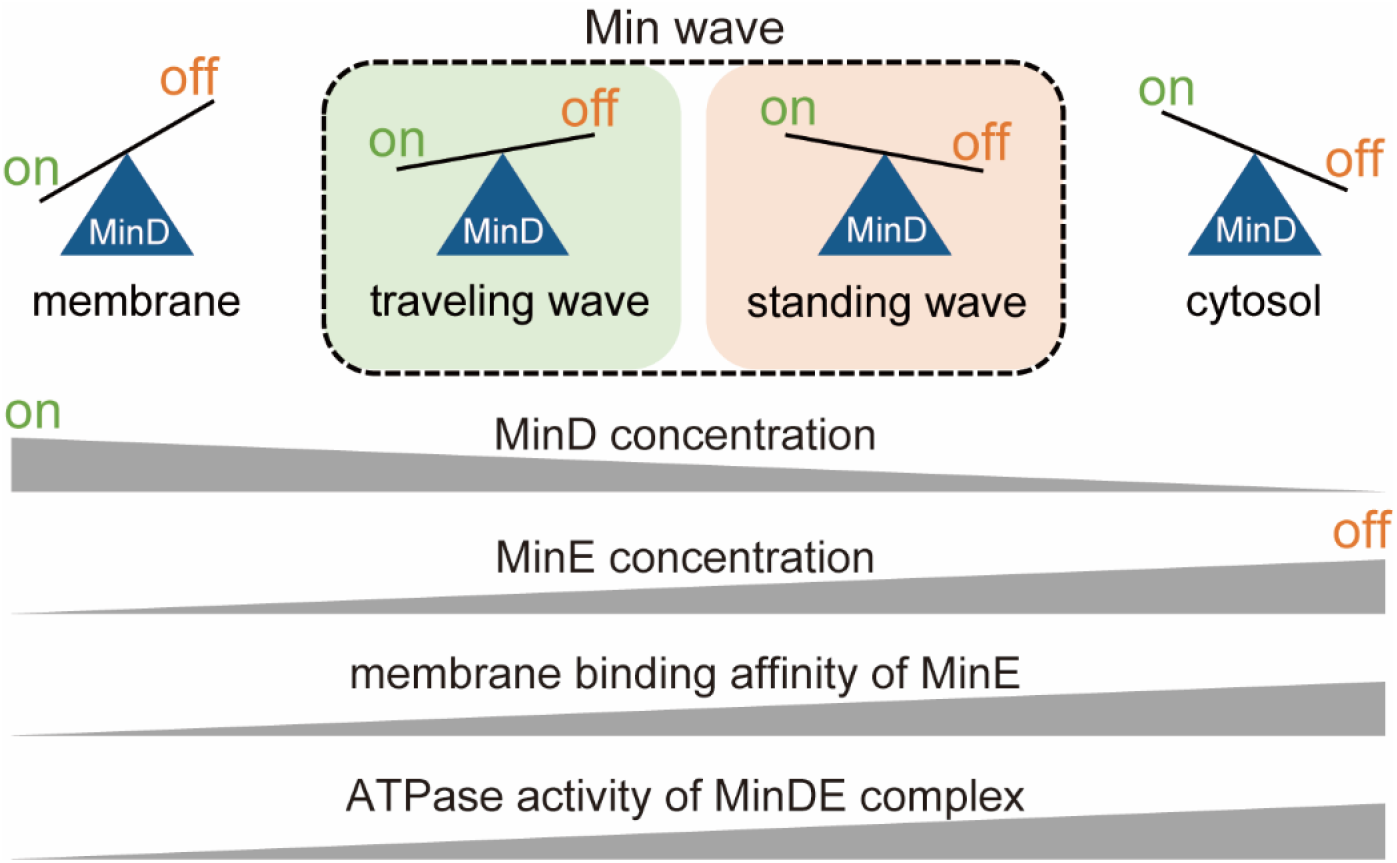
Summary of the results in this study. “on” and “off” denote the membrane binding and dissociation from the membrane of MinD, respectively. Parameters involved in membrane binding and dissociation from the membrane of MinD and its regulation of the spatiotemporal pattern generated by Min proteins are shown.

In the reconstitution system, both directions of spontaneous mode transition were observed: from the standing wave to the traveling wave and vice versa(*27, 28*). The present study shows that this is because the dominant Min wave mode is sensitive to the Min protein. However, this sensitivity is not preferable for living cells, which requires a standing wave for normal cell division under various external environments. In fact, a standing wave is observed in living cells even if the temperature is changed, although the period of the wave changes depending on the temperature(*47*). Geometry might be the reason the Min wave mode is not sensitive to parameters in living cells, unlike the reconstitution system, as previously proposed(*22, 23*). Min waves reconstituted in nonspherical artificial cells will provide an answer to this point in the future.

In this study, we also elucidated that the Min wave mode is controllable, and based on the findings, we achieved wave mode switching in artificial cells (Movies S7 and S8). Although the Min wave mode selection finding indicates that the process is deterministic, both experimental and theoretical results of cooling to the original temperature after increasing the temperature (Figs. 3C and 5E) and the dynamic increase in MinE concentration (Figs. 4B and 5F) suggest that the Min wave mode has hysteresis characteristics. The hysteresis is due to the non-linearity of the RDc systems, and its underlying mechanism is not clarified in the present study. A possible explanation of the origin of hysteresis is the multistability of the Min wave mode, as reported in previous studies(*25, 50, 51*). However, further analysis is needed to reveal the physical mechanisms underlying the phenomenon. Elucidation of the physical mechanism will provide a novel paradigm of the pattern formation theory for RDc and develop methods for further precise control of the wave modes.

Controlling Min wave mode selection can be used to expand the use and varieties of artificial cells. The time-averaged distribution was different between the two wave modes. The traveling wave generates a homogenous time-averaged distribution of MinD on the membrane, while the standing wave generates a minimum time average concentration of MinD at the center of living cells. In living cells, the cell division plane is determined by the difference in the time-averaged distribution. This is because MinC, which inhibits FtsZ polymerization for cell division initiation, does not localize at the minimum, and cell division starts at the center of the cell(*11*). Therefore, finding stable generation conditions for the standing wave will contribute to the reconstitution of bacterial cell division machinery in artificial cells. Moreover, because these two RDc wave modes are linked to cellular functions, such as motility (*6*)and cell division(*9, 10*), the transition between the traveling and standing wave modes of Min waves could be utilized as a cue to the switching dynamics of artificial cells in response to environmental signals. Hence, the findings of this study pave the way for the construction of molecular robots with flexible function switches using the mode transition of RDc waves.

Finally, we discuss the physiological significance of the elucidation of the mechanism Min wave mode selection in artificial cells. We showed not only the geometry, but also the parameters involved in the mode selection of Min waves. This finding explains why both standing and traveling waves have been observed in rod-shaped and spherical cells. Furthermore, the findings will lead to an understanding of the mechanism of spatiotemporal pattern formation using RDcs other than Min waves. For example, in mast cells, the traveling wave of actin is the dominant wave mode, but the standing oscillation of actin also appears due to the oscillation of PI(4,5)P_2_, which accompanies the oscillation of cytosolic calcium(*52*). Changes in parameters may influence the generation of this standing oscillation of actin waves, similar to the Min waves. As another example, our findings predict that standing waves can appear in the traveling wave of Rho and actin waves, in which no standing waves have been observed so far. To address these points in the future, we will illuminate novel roles for RDc waves in living cells.

## Materials and Methods

### Expression and purification of Min proteins

*Escherichia coli* BL21-CodonPlus(DE3)-RIPL cells (Agilent Technologies, Santa Clara, CA, USA) were transformed with either pET15-MinD, pET29-MinE, or pET15-msfGFP-MinC and cultivated in LB medium with ampicillin (MinD and msfGFP-MinC) or kanamycin (MinE) at 37°C. The proteins were expressed by 1 mM IPTG at OD_600_ = 0.8 (MinD) or at OD_600_ = 0.2 (MinE and msfGFP-MinC) and the cells were further cultivated for 1.5 h (MinD) or for 3-4 h (MinE and msfGFP-MinC), respectively. The cells were collected by centrifugation at 8000 × g for 2 min at 4°C, resuspended in the lysis buffer [50 mM Tris-HCl (pH7.6), 300 mM NaCl,1 mM PMSF, 10 mM imidazole, and 1 mM DTT], and sonicated using Sonifier250 (Branson, Danbury, CT, USA). In the case of His-MinD, imidazole concentration of the lysis buffer was 20 mM and 0.1 mM ADP was added to the lysis buffer. The soluble fraction in the lysate was fractionated by centrifugation at 20000 × g for 30 min at 4°C. The supernatant was filtered by HPF Millex HV (Merck Millipore, Billerica, MA, USA) and mixed with 1 mL Ni Sepharose 6Fast Flow (Cytiva, Tokyo Japan) (MinD) or cOmplete His-Tag Purification Resin (Roche, Basel, Switzerland) (MinE and msfGFP-MinC). After shaking for 30 min at 4°C, the mixture was loaded to a Poly-Prep Chromatography Column (Bio-Rad, Hercules, CA, USA) and washed by adding 25 mL wash buffer [50 mM Tris-HCl (pH7.6),300 mM NaCl,1 mM PMSF, 0.1 mM EDTA, 20 or 25 mM imidazole, and 10% Glycerol]. Then, His-tagged proteins were eluted with 2 mL elution buffer [50 mM Tris-HCl (pH7.6),300 mM NaCl,1 mM PMSF, 0.1 mM EDTA, 250 or 500 mM imidazole, and 10% Glycerol]. The elution buffer was exchanged to the storage buffer [50 mM HEPES-KOH (pH7.6), 150 mM GluK,0.1 mM EDTA, 10% Glycerol, 0.1 mM ADP in the case of MinD] by repeating the ultrafiltration using an AmiconUltra-4 3K (MinE) or 10K (MinD and msfGFP-MinC) (Merk Millipore) and the addition of the storage buffer. The proteins were separated by SDS-PAGE and the concentration was estimated by quantifying the band intensity after CBB staining using Fiji software (National Institutes of Health, Bethesda, MD, USA).

### Self-organization assay inside microdroplets covered with lipids

25 mg/mL *E*.*coli* polar lipid in chloroform (Avanti, Alabaster, AL, USA) was dried by gentle Argon gas flow and dissolved in mineral oil (Nacalai Tesque, Kyoto, Japan) to a lipid concentration of 1 mg/mL in glass tubes. The lipid mixture was sonicated for 90 min at 60°C using Bransonic (Branson), and mixed by vortexing for 1 min.

In the case of the standard condition, the inner solution consisted of 0.1 μM His-msfGFP-MinC, 1 μM His-MinD, 1 μM MinE-His, 2.5 mM ATP and 100 mg/mL BSA in the reaction buffer [25 mM Tris-HCl (pH 7.6), 150 mM GluK, 5 mM GluMg]. Prior to preparation of the mixture, BSA (A6003, Sigma-Aldrich, St. Louis, MO, USA) was dissolved in water and washed with the reaction buffer by using AmiconUltra-0.5 50K (Merck Millipore), and the concentration was quantified by Pierce BCA Protein Assay kit (Thermo Fisher Scientific, Waltham, MA, USA). The concentrations of each component were varied as indicated. 1 μL of the inner solution was added to the 50 μL lipid mixture in microtubes, and the microdroplets were obtained by tapping microtubes. 20 μL of the droplets mixture was placed into two glass coverslip slits. The self-organization of Min proteins inside the droplets was observed using fluorescent microscope (AxioObserver Z1; Carl Zeiss, Jena, Germany). The time-lapse images obtained were processed by using Fiji software.

In the case of the investigation of the effect of temperature, a glass heating stage (Tpi-SQFTX, Tokai Hit, Shizuoka, Japan) was set on the microscope, and the prepared slide was placed on the glass heating stage immediately after sample preparation.

In the case of the observation of the early stage of Min wave emergence, 10 mM creatine phosphate, 0.1 μM creatine kinase and 2 mM ADP were added to the inner solution instead of ATP. After making the prepared slide of microdroplets mixture, the self-organization of Min proteins was observed immediately.

### Protein synthesis by the PURE system in microdroplets and in test tubes

PUREfrex ver. 1.0 (Gene Frontier, Chiba, Japan) was used to synthesize proteins. For the investigation of the regulation of the amount of protein synthesized, the mixture of small molecules [20 mM HEPES-KOH (pH7.6), 30 ng/μL tRNA, 10 μg/mL 10-formyltetrahydrofolate, 20 mM creatine phosphate, 2 mM dithiothreitol (DTT), 180 mM potassium glutamate, 2 mM spermidine, and 14 mM magnesium acetate at final], 20 amino acids mixture, NTPs mixture [2 mM ATP, 2 mM GTP, 1 mM CTP and 1 mM UTP at final], 1 nM DNA encoding NanoLuc at final were added to the solution II and III of PUREfrex ver. 1.0 in the test tubes. The concentrations of 20 amino acids were varied as indicated. After the 4 h reaction at room temperature, the concentration of NanoLuc was quantified by Nano-Glo Luciferase Assay System (Promega, Madison, WI, USA). In the case of the self-organization assay with MinE synthesis inside microdroplets, the mixture of 0.1 μM His-msfGFP-MinC, 1 μM His-MinD, 0.7 μM MinE-His, 100 mg/mL BSA, small molecules described above, 20 amino acids mixture, NTPs mixture, 1 nM DNA encoding MinE and the solution II and III of PUREfrex ver. 1.0 was encapsulated in microdroplets. The microdroplets encapsulating the inner solution were obtained as described above and placed into two glass coverslip slits. The change of the dynamics of self-organized Min proteins during incubation was observed using fluorescent microscope.

### ATPase assay

For the investigation of the temperature dependence of ATPase activity of MinD, ATPase activity of MinD at various temperatures was tracked by measuring the increase in inorganic phosphate (Pi) concentration. To prepare small unilamellar vesicles (SUVs) used as lipids, 25 mg/mL *E*.*coli* polar lipid in chloroform was dried under gentle Argon gas flow. The lipid film was further dried in a desiccator for 3 h at room temperature. The reaction buffer for the Min wave observation was added to the lipid film to adjust the lipid concentration of 4 mg/mL and incubated at room temperature overnight. The lipid solution was vortexed for 1 min and extruded by using Avanti Mini Extruder (Avanti). In the extrusion process, the lipid solution was passed through polycarbonate membranes with pore size 1.0 μm, 0.4 μm, and 0.05 μm, in this order, to obtain the SUVs. The reaction mixture for ATPase assay consisted of 1 μM His-MinD, 1 μM MinE-His, 2.5 mM ATP, and 1 mg/mL SUVs solution in the reaction buffer. The mixture was incubated at 25°C, 29°C, and 37°C for 0-30 min after 5 min incubation at 25°C. The concentration of Pi of each reaction mixture incubated under three temperature conditions from 0 min to 30 min was quantified by using BIOMOL Green (Enzo Life Science, Farmingdale, NY, USA). ATPase rates were calculated from the slope of the increase in Pi with time.

### Theoretical model

We performed numerical simulations of the model based on the previous works(*28, 43, 49*). ATP-MinD, ADP-MinD, and MinE concentrations inside a spherical membrane with its radius, *R*, were denoted by *c*_*DT*_, *c*_*DD*_, and *c*_*E*_, respectively. Concentrations of MinD, MinE, and their complex (MinDE) bound to a membrane were denoted by *c*_*d*_, *c*_*de*_, and *c*_*e*_, respectively. Each reaction shows a rate, *ω*, specified by its subscript. Recruitment of MinD onto the membrane from the cytosol is given by *ω*_*D*_ and *ω*_*dD*_, where in the latter reaction, membrane-bound MinD recruits MinD in the cytosol. Recruitment of MinE in the cytosol by membrane-bound MinD is described by *ω*_*E*_. The binding rate *ω*_*ed*_ of MinD and MinE on the membrane is assumed to be fast. Attachment and detachment of MinD from/to the cytosol are described by *ω*_*eE*_ and *ω*_*e*_, respectively. We assumed the same diffusion constants for proteins bound to the membrane, and set them to be unity without loss of generality. We also assumed that the bulk diffusion constants of unbound proteins are the same, and are denoted by *D*. The time scale was normalized by 1/*ω*_*e*_. The model is given by the following equations:

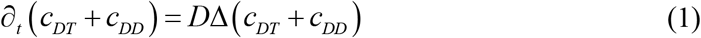

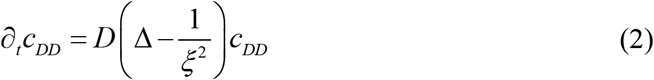

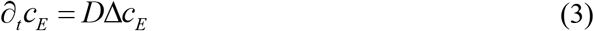

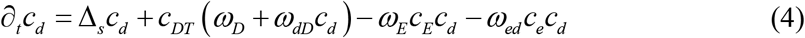

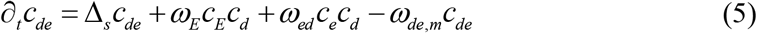

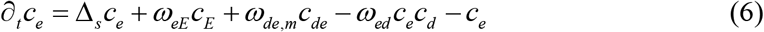

Here, Δ and Δ_*s*_ denote the Laplacian operator in three-dimensional bulk space and the Laplace-Bertrami operator on the two-dimensional surface, respectively. The boundary conditions of Eqs.(1) are

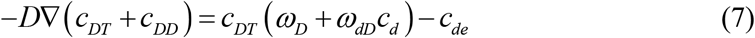

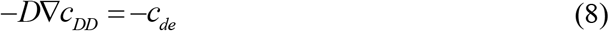

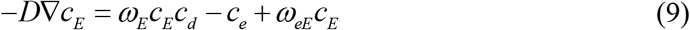

The length scale associated with ATP hydrolyzation is denoted by 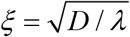 where *λ* is the rate of ATP hydrolyzation. We set *λ* =1. Eqs. (1)-(5) are solved using the software of the Finite Element method, COMSOL. To compute wave generation in a broad range of parameters, we used a two-dimensional system in which the cytosol is inside the disk, and the membrane is on a circle. We have confirmed that the same results are obtained at the selected data points in three dimensional systems. We mainly chose the parameters as *D* = 100, *ω*_*D*_ = 0.1, *ω*_*dD*_ = 5.0, *ω*_*E*_ = 0.1, *ω*_*ed*_ = 100, *ω*_*de,m*_ = 1.0, and *ω*_*eE*_ = 0.04 (*28*), and varied *ω*_*dD*_, *ω*_*E*_, and, *ω*_*de,m*_ in the text. In the parameter space of *ω*_*E*_ − *ω*_*dD*_, each parameter was discretized in 21 mesh points. We also used the same data points in the parameter space of MinE-MinD. For each parameter set, we perform the simulations for the time *T* = 2000. The initial condition was chosen to be the Gaussian random distribution in the cytosol. The wave generation occurred around *t* = 100 as a standing wave, and then the wave switched to the traveling wave when the parameters are at its state. After the relaxation time *T* ≥1000, the wave states were estimated.

Inhomogeneity of the concentration field was expressed by the amplitude of each mode in the expansion of the concentration *c*_*d*_ by the Fourier expansion in the polar coordinates (*r,θ*) as

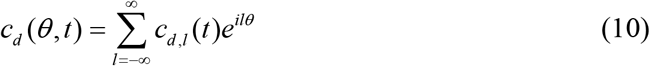

For example, the uniform distribution of MinD on a membrane was expressed by the l=0 mode and its norm, |*C*_*d*_, 0 |, whereas the first mode (l = 1) corresponds to the inhomogeneous concentration field of a single wave, which is characterized by the norm 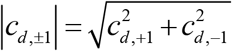. The state diagram was drawn by the following criteria: The wave state is at ⟨|*c*_*d*, ±1_(*t*)|⟩ ≥ 0.1 and max |*c*_*d*, ±1_ (*t*)|− min|*c*_*d*, ±1_ (*t*)|≥ 0.1, where ⟨ · ⟩ denotes the average over time. The traveling and standing waves were detected by the dynamics of the phase *ϕ*(*t*) = tan^−1^ *c*_*d*, −1_ / *c*_*d*, 1_ of MinD on the membrane. When max *θ* (*t*) − min *θ* (*t*) ≥ 3*π* / 2, the wave is traveling wave, whereas the standing wave is at max *θ* (*t*) − min *θ* (*t*) < 3*π* / 2.

When we varied the parameter *ω*_*de,m*_, we applied the stepwise change from *t* = 500 to *t* = 1000, that is, *ω*_*de,m*_ is larger at 500 ≤ *t* ≤1000, while *ω*_*de,m*_ = 1 otherwise. The parameter change was performed continuously with a time width of 1. When we varied the concentration of MinE, we added the source term in the right-hand side of Eq.(3). From *t* = 500, we added 0.001 at each time until the total MinE became the target value (either MinE=1.0 or MinE=1.5). This procedure implies that the MinE concentration in the cytosol changes linearly in time.

## Acknowledgments

We thank Prof. Miho Yanagisawa (The University of Tokyo) for helpful discussion and an advice on the manuscript.

## Author contributions

Conceptualization: KF

Methodology: ST, NY, KF

Formal analysis: ST, NY, KF

Funding acquisition: NY, KF

Investigation: ST, NY, KF

Visualization: ST, NY

Project administration: KF

Supervision: NY, ND, KF

Writing—original draft: ST, NY, KF

Writing—review & editing: ST, NY, ND, KF

## Data and materials availability

All data are available in the main text or the supplementary materials.

## Supplementary Materials

This PDF file includes:

Figs. S1 to S7

Movie S1 to S10

