## Supplementary Information for "Mode selection mechanism in traveling and standing waves revealed by Min wave reconstituted in artificial cells"

**This PDF file includes:**

Figs. S1 to S7
Movies S1 to S10

**Other Supplementary Materials for this manuscript include the following:**
Movies S1 to S10

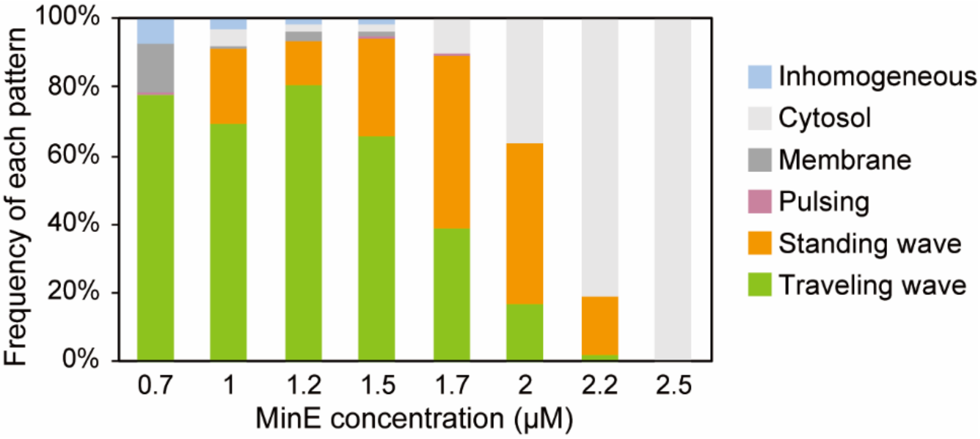


**Fig. S1.**

Supplementary Figure S1. Frequencies of each pattern generated by the Min proteins in artificial cells at various MinE concentrations (n = 160-217).

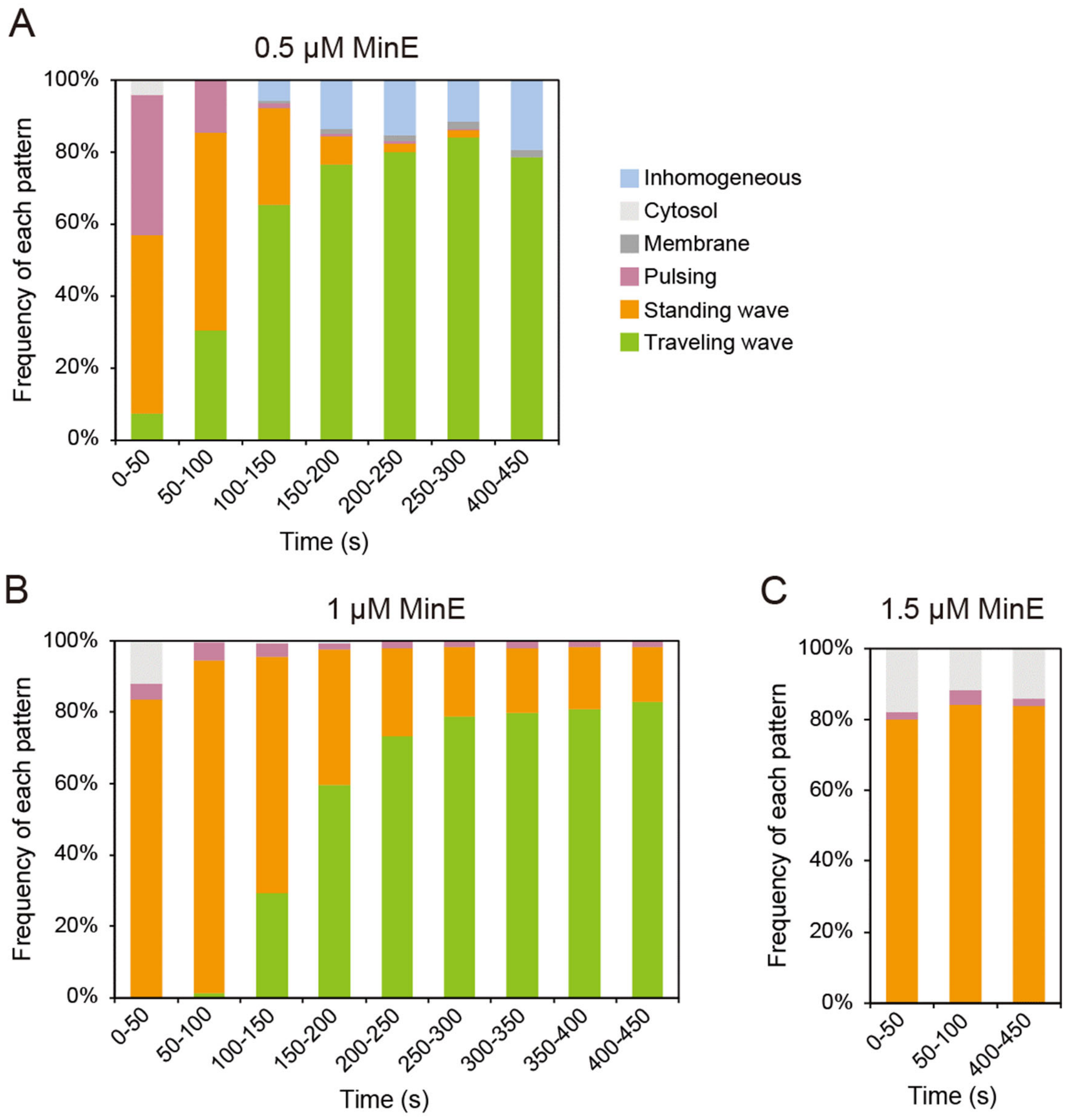


**Fig. S2.**

Time development of the frequencies of each pattern generated by the Min system in artificial cells at (**A**) 0.5 (n = 150-165), (**B**) 1.0 (n = 218-332), and (**C**) 1.5 µM MinE (n = 90-99).

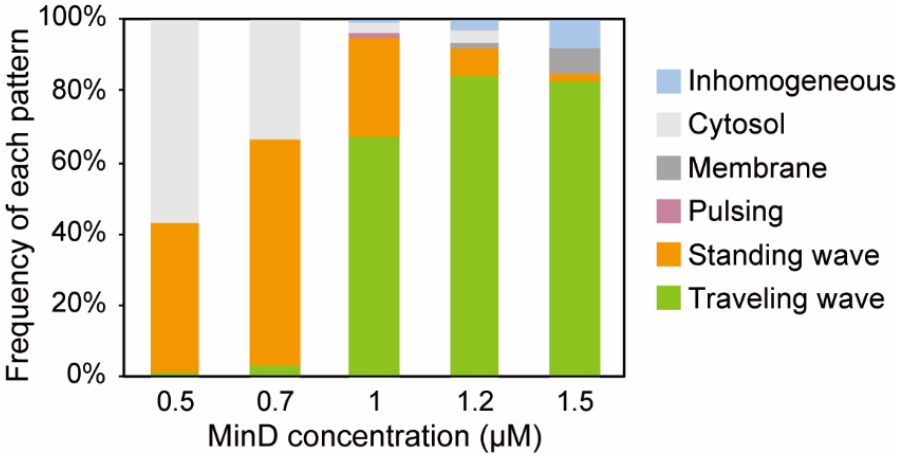


**Fig. S3.**
Frequencies of each pattern generated by the Min proteins in artificial cells at various MinD concentrations (n = 158-249).

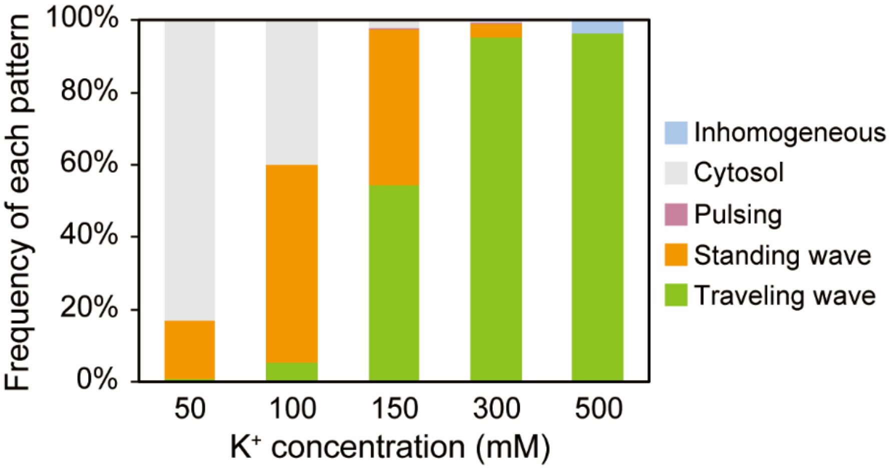


**Fig. S4.**

Frequencies of each pattern generated by the Min system in artificial cells at various K+ concentration (n = 130-213).

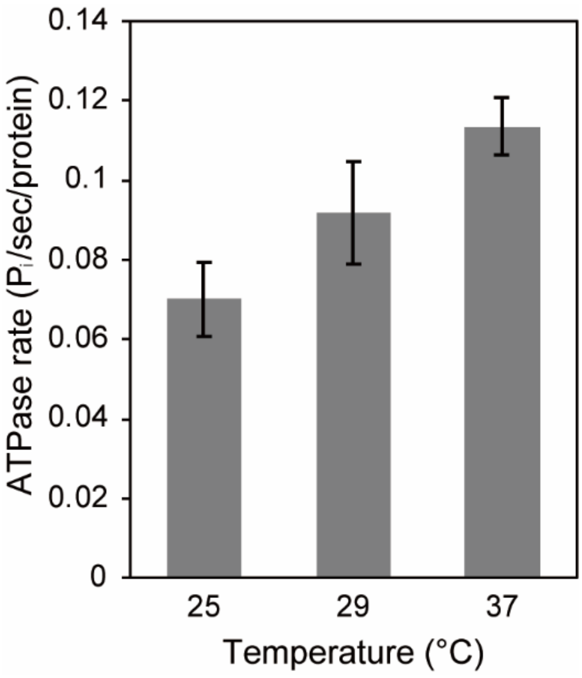


**Fig. S5.**
ATPase rates of MinDE complex with increasing temperature. Mean ± standard error (n = 4) are shown.

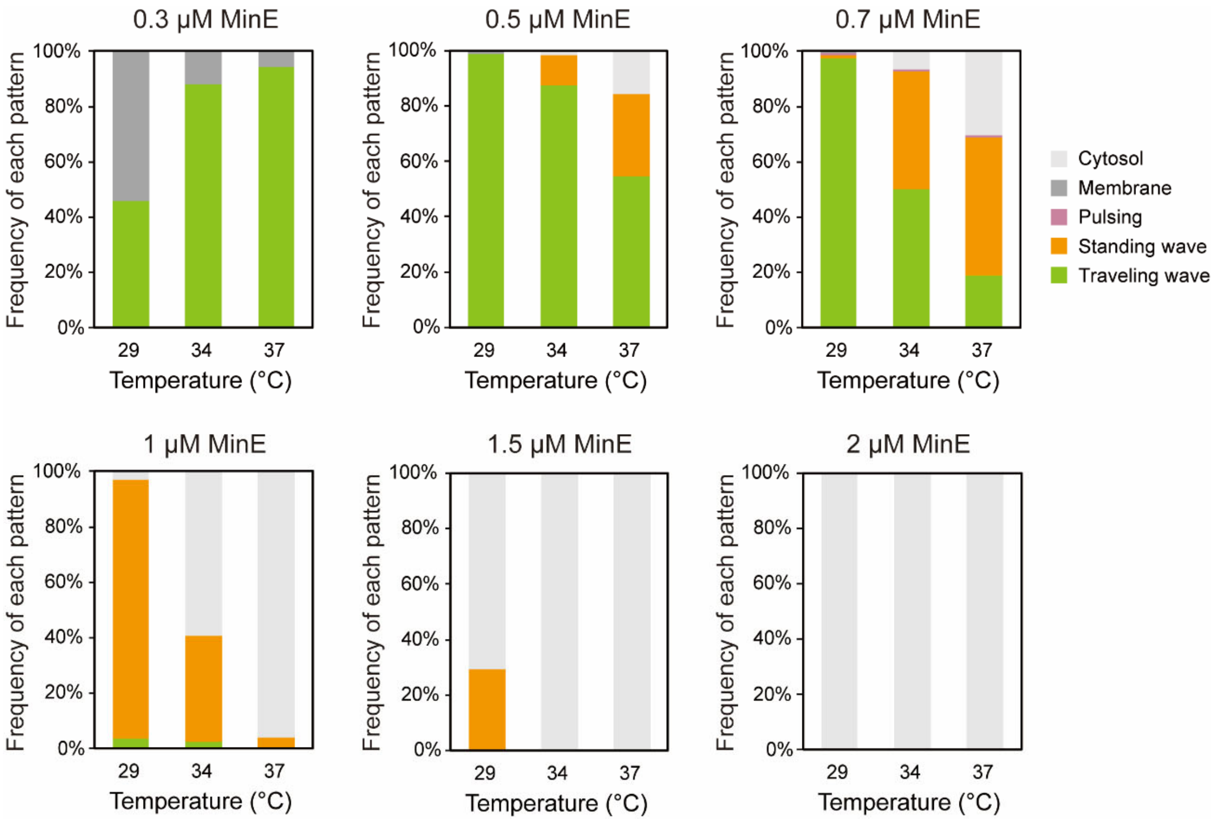


**Fig. S6.**
Frequencies of each pattern generated by the Min system in artificial cells at various MinE concentrations at 29, 34, 37°C (n = 88-162, 0.7 μM MinE at 37°C: n = 280).

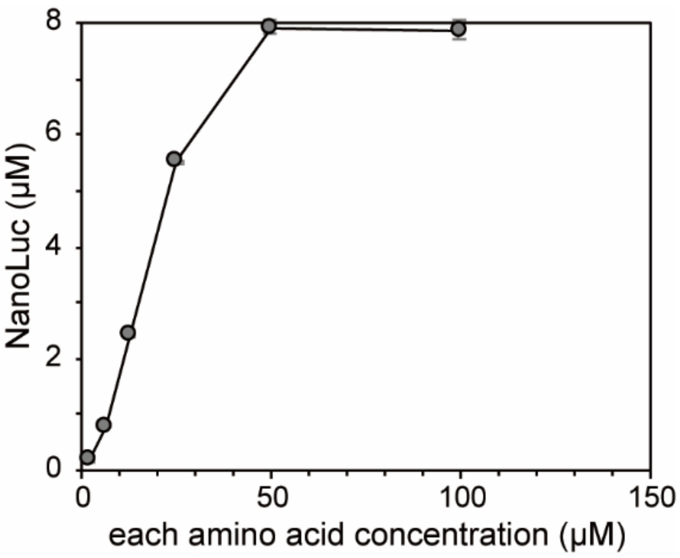


**Fig. S7.**

The amount of NanoLuc synthesized by PURE system depends on each amino acid concentration. Mean ± standard deviation (n = 3) are shown.

**Movie S1.**
Min wave in artificial cells with different MinE concentrations (scale bar: 10 μm)

**Movie S2.**
Initial Min wave mode transition at 0.5 μM MinE

**Movie S3.**
Initial Min wave mode transition at 1.0 μM MinE

**Movie S4.**
Initial Min wave mode transition at 1.5 μM MinE

**Movie S5.**
Min wave in artificial cells with different MinD concentrations (scale bar: 10 μm)

**Movie S6.**
Min wave in artificial cells with different potassium concentrations (scale bar: 10 μm)

**Movie S7.**
Min wave mode transition by temperature shift (scale bar: 10 μm)

**Movie S8.**
Min wave mode transitions by temperature shifts (scale bar: 10 μm)

**Movie S9.**
Min waves in artificial cells with MinE synthesis (1.5 μM aa)

**Movie S10**
Min waves in artificial cells with MinE synthesis (2 μM aa)
